# Cas9 targeted nanopore sequencing with enhanced variant calling improves *CYP2D6*-*CYP2D7* hybrid allele genotyping

**DOI:** 10.1101/2022.03.31.486504

**Authors:** Rubben Kaat, Tilleman Laurentijn, Deserranno Koen, Tytgat Olivier, Deforce Dieter, Filip Van Nieuwerburgh

**Author notes:** equal contribution (Rubben Kaat, Tilleman Laurentijn). (KR). (LT). (KD). (OT). (DD). (FV).

## Abstract

*CYP2D6* is one of the most challenging pharmacogenes to genotype due to the high similarity with its neighboring pseudogenes and the frequent occurrence of *CYP2D6-CYP2D7* hybrids. Unfortunately, most current genotyping methods are therefore not able to correctly determine the complete *CYP2D6-CYP2D7* sequence. Therefore, we developed a genotyping assay to generate complete allele-specific consensus sequences of complex regions by optimizing the PCR-free nanopore Cas9-targeted sequencing (nCATS) method combined with adaptive sequencing, and developing a new comprehensive long read genotyping (CoLoRGen) pipeline. The CoLoRGen pipeline first generates consensus sequences of both alleles and subsequently determines both large structural and small variants to ultimately assign the correct star-alleles. In reference samples, our genotyping assay confirms the presence of *CYP2D6-CYP2D7* large structural variants, single nucleotide variants (SNVs), and small insertions and deletions (INDELs) that go undetected by most current assays. Moreover, our results provide direct evidence that the *CYP2D6* genotype of the NA12878 DNA should be updated to include the *CYP2D6-CYP2D7* *68 hybrid and several additional single nucleotide variants compared to existing references. Ultimately, the nCATS-CoLoRGen genotyping assay additionally allows for more accurate gene function predictions by enabling the possibility to detect and phase *de novo* mutations in addition to known large structural and small variants.

**Author Summary:** During the last decades, the usefulness of personalized medicine has become increasingly apparent. Directly linked to that is the need for accurate genotyping assays to determine the pharmacogenetic profile of patients. Continuing research has led to the development of genotyping assays that perform quite robustly. However, complex genes remain an issue when it comes to determining the complete sequence correctly. An example of such a complex but very important pharmacogene is *CYP2D6*. Therefore, we developed a genotyping assay in an attempt to generate complete allele-specific consensus sequences of *CYP2D6*, by optimizing a targeted amplification-free long-read sequencing method and developing a new analysis pipeline. In reference samples, we showed that our genotyping assay performed accurately and confirmed the presence of variants that go undetected by most current assays. However, the implementation of this assay in practice is still hampered as the selected enrichment strategies inherently lead to a low percentage of on-target reads, resulting in low on-target sequencing depths. Further optimization and validation of the assay is thus needed, but definitely worth considering for follow-up research as we already demonstrated the added value for generating more complete genotypes, which on its turn will result in more accurate gene function predictions.

## Introduction

Genotyping is one of the most important aspects of personalized medicine, particularly within the context of pharmacogenetics (1,2). In many medical disciplines, pharmacogenetic genotyping is used to predict a patient’s phenotype in order to adjust therapy (3,4). Especially the genetic variation in drug-metabolizing enzymes significantly contributes to the differing benefit-risk balance of certain drugs between patients (1,4). One of the essential drug-metabolizing enzymes is Cytochrome P450 2D6 (CYP2D6), as it is responsible for the metabolization or bioactivation of 20 to 30% of the clinically used drugs (4). Therefore, accurate genotyping assays for this gene are of major importance. However, although *CYP2D6* is a relatively small gene spanning only 4400 nucleotides, accurate genotyping of this gene is challenging. First of all, the *CYP2D6* gene is surrounded by two pseudogenes showing 94% sequence similarity with *CYP2D6*, which complicates the genotyping of this gene. Furthermore, *CYP2D6* is one of the most polymorphic human genes, with over 100 star(*)-alleles and over 400 sub-alleles (5,6). This star- and sub-allele nomenclature does not only encompass small sequence variations, such as single nucleotide variants (SNVs) or insertions and deletions smaller than 50 bp (INDELs), but also large structural variants, such as gene deletions and multiplications. On top of that, the possible formation of hybrids with its nearest pseudogene *CYP2D7* poses an additional major challenge when a comprehensive genotype is desired (5–8).

In addition to the gene structure, a second important factor for accurate genotyping is the applied genotyping assay. Various assays have been used for genotyping the *CYP2D6* gene, such as polymerase chain reaction (PCR), microarrays, or short-read (SR) next-generation sequencing (NGS) (9–11). However, most currently used assays target only a limited subset of pre-selected SNVs (12–14). Only a few assays determine the correct genotype based on multiple detected SNVs and copy number variations in each allele (13,15,16). Nevertheless, as 35.4% of the variant-drug interactions described in the Clinical Annotations of PharmGKB are based on complete alleles containing all its variants, more comprehensive genotyping assays could be valuable in the clinical practice (7,13,17). SR NGS technologies can identify most individual variants in a genome, but mapping short reads to homologous elements, such as those in *CYP2D6* and *CYP2D7*, is error-prone. On top of that, phasing of short-read data is not straightforward, as it typically requires supplemental statistical phasing based on known allele structures in the population or parental genotypic data (18).

Recently, efforts have been realized to comprehensively genotype *CYP2D6* in an attempt to overcome these mapping and phasing problems (18–22). Different studies have shown that long-read sequencing platforms can discover new variants and determine the correct allele structure (19,20). However, these studies use long-range PCR to capture the targeted region, which is prone to template switching. This, on its turn, results in chimeric PCR products and introduces phasing errors (23). To avoid the application of long-range PCR (LR-PCR), a new enrichment strategy, called nanopore Cas9-targeted sequencing (nCATS), was introduced by Gilpatrick *et al*. (24). This strategy uses targeted cleavage of DNA with Cas9, followed by selectively ligating adapters for nanopore sequencing. However, ligation of nanopore adapters to random breakage points also generates a considerable number of so-called background reads, bringing the percentage of on-target reads down to merely 0.5% to 15% of the sequenced reads in practice (24–26). To increase the number of reads on-target, a second PCR-free enrichment strategy for nanopore sequencing, called adaptive sequencing (AS), could be used in addition. AS refers to the ability of a nanopore sequencer to reject individual molecules in real-time while they are being sequenced, and as such, does not involve additional steps in the library preparation (27).

The aim of this study was to develop a new assay for correct and complete genotyping of complex regions such as the *CYP2D6* gene. This genotyping assay consists of two important steps that need to be optimized. The first step entails the generation of long reads using a PCR-free enrichment strategy combined with nanopore sequencing. Therefore, the nCATS and combined nCATS-AS enrichment strategies were both tested on the *CYP2D6-CYP2D7* locus. For this purpose, a guide RNA (gRNA) panel was optimized to enrich *CYP2D6* and *CYP2D7* from human DNA samples. The second step aims to correctly elucidate both large structural and small variants to determine the alleles of cell lines that might contain both types of variants. However, the currently existing tools do not combine the detection of large structural and small variants in one pipeline (28–31). Consequently, smaller variants cannot be detected in regions with large structural variants, and large structural variants are not taken into account when small variants are detected with currently available tools. This might lead to the incorrect determination of gene sequences and complicate the correct assignment of star-alleles. Therefore, we developed a new comprehensive long read genotyping (CoLoRGen) pipeline that is able to simultaneously detect both large structural and small variants in complex genes such as *CYP2D6*.

## Materials and methods

### Cell cultivation, DNA extraction, and nCATS

Two lymphoblast cell lines, HG01990 and GM19785, of which the *CYP2D6* genotype is well-known in the literature (15,16), were cultivated and subsequently subjected to DNA extraction to obtain the samples for the experiments conducted within this study. Cells were washed every three to four days to an optimal cell density for successful cell growth of 300.000 cells/mL. The old medium was washed away through 5-minute centrifugation at 500 to 600g, after which a new medium was added. The medium contained 1% penicillin-streptomycin, 15% fetal bovine serum, and 2mM L-glutamine in Roswell Park Memorial Institute (RPMI) 1640 medium. DNA samples were extracted using the Dneasy Blood & Tissue kit (Qiagen, Venlo, The Netherlands), quantified using the Qubit fluorometer with the dsDNA High Sensitivity Assay kit (ThermoFisher Scientific, Waltham, MA, USA), and stored at 4°C until further processing. A Zymo DNA Clean & Concentrator purification step (Zymo Research, Irvine, CA, USA) was performed to remove the excess salts, whereby the DNA was eluted in water. The length of the eluted DNA fragments was measured on a Femto Pulse using the Agilent Genomic DNA 165 kb kit (Agilent Technologies, Santa Clara, CA, USA) according to the manufacturer’s recommendations.

The library preparation of the samples was performed according to the ‘Cas9 targeted sequencing’ Oxford Nanopore Technologies (ONT) protocol, using the LSK-110 kit (ONT, Oxford, UK) (Figure 1). Nine guide RNAs (gRNAs) were designed with the CHOPCHOP tool (32). Four of them were designed to cut upstream *CYP2D6*, two downstream *CYP2D7*, and three between *CYP2D6* and *CYP2D7* (Table S1). The gRNAs cutting between the two genes were added to ensure sufficient depth on *CYP2D6* for reliable variant calling. The efficiency of the gRNAs was assessed beforehand in preliminary sequencing runs using purchased NA12878 DNA. After selecting the seven most efficient gRNAs, two separate gRNA pools were created. As shown in Figure 1, pool A only contained seven gRNAs that cut upstream *CYP2D6* or downstream *CYP2D7*, whereas pool B also contained a gRNA that hybridizes between the two genes. The use of two separate pools, one without gRNAs that cut between the genes, is necessary to obtain reads covering the complete *CYP2D6-CYP2D7* locus. Active RNA ribonucleoprotein complex (RNP) complexes were subsequently created in two separate tubes, using Alt-R® *S. pyogenes* HiFi Cas9 nuclease V3 (IDT, Leuven, Belgium), *S. pyogenes* Cas9 tracrRNA (IDT, Leuven, Belgium), and one of the pools with *S. pyogenes* Cas9 Alt-R(tm) gRNAs (IDT, Leuven, Belgium).

**Figure 1.**
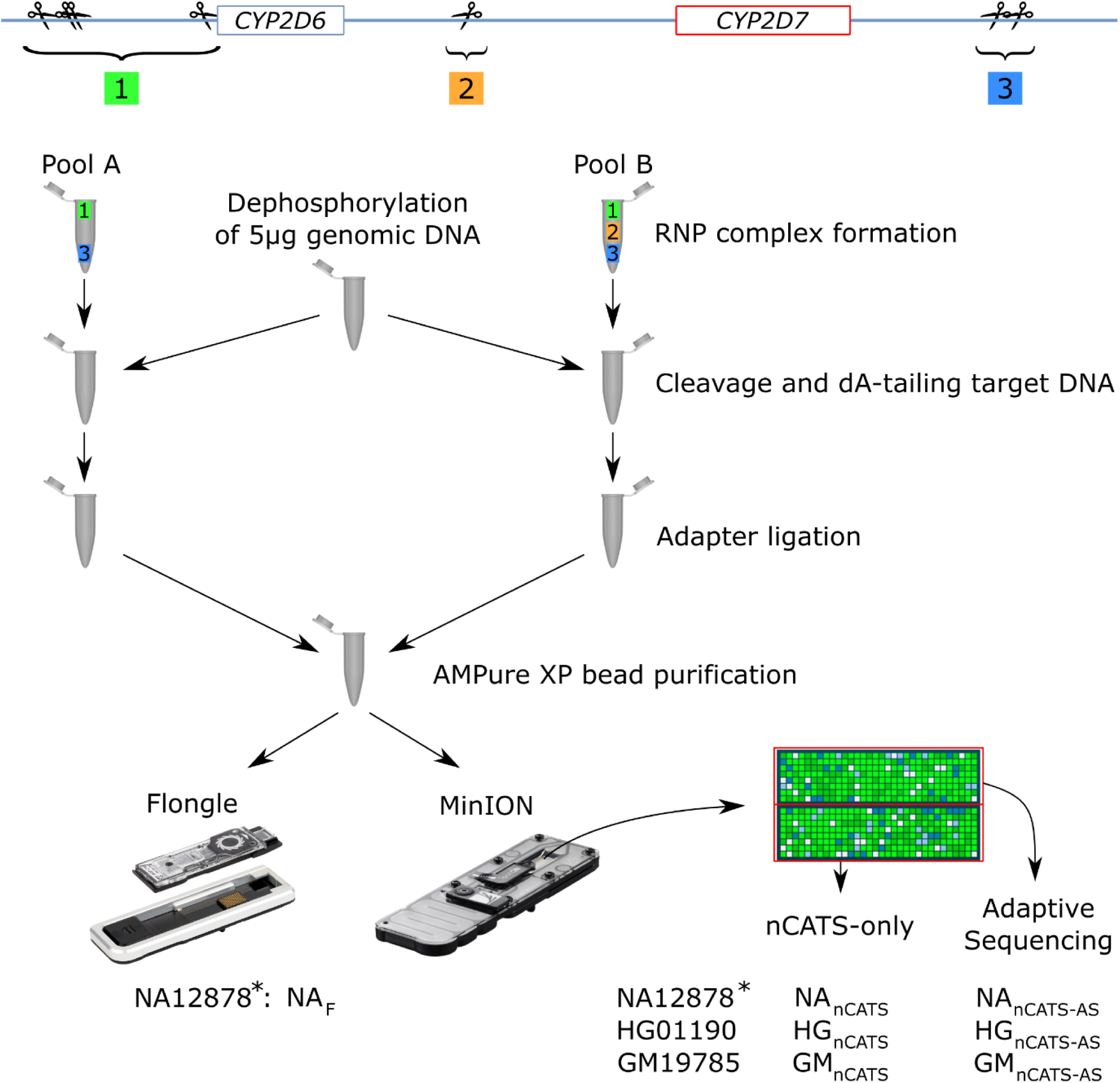
Enrichment and sequencing workflow adapted from the ‘Cas9 targeted sequencing’ protocol from ONT. Two different pools of gRNAs were made. Pool A only contains gRNAs that cut upstream and downstream the *CYP2D6-CYP2D7* locus, Pool B also contains a gRNA that cuts between *CYP2D6* and *CYP2D7*. After dephosphorylation of the genomic DNA, half of the DNA was cleaved by the RNP with the gRNAs of Pool A, and the other half was cleaved by the RNP with the gRNAs of Pool B. After cleavage, the adaptors were ligated at the cleavage site. Next, the two pools were mixed again and purified with AMPure XP beads. The NA12878 libraries were sequenced on a Flongle (NA_F_) and on a MinION flow cell. The HG01190 and GM19785 libraries were only sequenced on a MinION flow cell. On the runs using a MinION flow cell, half of the pores were controlled by the adaptive sequencing software (NA_nCATS-AS_, HG_nCATS-AS_, and GM_nCATS-AS_), and the other half sequenced conventionally (NA_nCATS_, HG_nCATS_, and GM_nCATS_). *: The NA12878 libraries were used for preliminary optimization purposes and were created with only one pool containing 8 (NA_F_) or 9 gRNAs (NA_nCATS-AS_ and NA_nCATS_).

Five µg of purchased NA12878, extracted HG01990, and extracted GM19785 DNA was dephosphorylated using Quick Calf Intestinal Phosphatase (NEB, Ipswich, MA, USA). The dephosphorylated NA12878 DNA was added to one RNP complex pool with 9 and 8 gRNAs for the MinION and Flongle library, respectively. The dephosphorylated DNA from the HG01990 and GM19785 cell lines was equally divided between the two Cas9 RNP complex pools. Subsequently, the target DNA was cleaved by the active RNP complex, and Taq Polymerase (NEB, Ipswich, MA, USA) was added for dA-tailing. Next, adapters were ligated to the newly produced DNA ends at the Cas9 cleavage sites by adding 5 µL of Adapter mix II, 20 µL of Ligation Buffer, and 10 µL NEBNext Quick T4 DNA Ligase (NEB, Ipswich, MA, USA) to the separate tubes. As the Cas9 enzyme remains bound to the DNA on the 5’-side of the cleavage site, adapters are preferentially ligated on the 3’-side of the cleavage site. After adapter ligation, the libraries were cleaned using a 0.3x volume of AMPure XP beads (Beckman Coulter, High Wycombe, UK). First, 80 µL TE of pH 8 (IDT, Leuven, Belgium) was added to each tube. For the HG01990 and GM19785 cell lines, the two separate tubes were pooled before adding the beads. 250 µL Long Fragment Buffer was subsequently used to wash the beads twice. After that, the beads were resuspended in 10 and 14 µl Elution Buffer during a 30-minute incubation at room temperature for the Flongle and MinION libraries, respectively. Before loading on a Flongle and MinION flow cell, 15 and 37.5 µL Sequencing Buffer, and 10 and 25.5 µL of Loading Beads were added to 5 and 12 µL of the eluate, respectively. The DNA libraries were sequenced using an R9.4 Flongle or MinION flow cell on a GridION device (ONT, Oxford, UK), and the AS software was activated on half of the pores of the MinION flow cells. The flow cells ran up to 48h to obtain the maximum number of reads possible and were controlled and monitored using the MinKNOW software.

### Data analysis, variant calling, and star-allele assignment

The raw sequencing data was basecalled using the high accuracy model of Guppy (v5.0.7). Raw reads were saved in fastq format, and only reads with a quality score above 8 were used for further analysis. These reads were subsequently split up into two groups, based on whether they were generated by pores controlled by the AS software or by pores that sequenced conventionally. All reads from the latter group were used for further data analysis, whereas only the positively selected reads from the first group were used in downstream analysis.

The data was processed with our in-house developed CoLoRGen pipeline to correctly assign both SNVs and INDELs as well as large structural variants in the basecalled data. To detect all these variants at once, several consecutive steps were carried out by the CoLoRGen pipeline (Figure 2). First, the reads were mapped against the human GRCh38 reference genome using Minimap (v2.18) (Figure 2A). Only the reads that mapped on the target region were retained for further analysis. Variant calling was performed on these reads using the Medaka Variant pipeline (v1.4.3). Based on the called SNVs and INDELs, the reads were split into two alleles using WhatsHap (v1.1). Breakpoints of large structural variants were defined for each allele separately, based on the starting points of clipping ends and the mapping coordinates of these clipping ends when mapped separately (red and green reads in Figure 2A, respectively). Only breakpoints covered by at least three reads were considered in order to obtain accurate structural variant calling. In the next step, an adjusted GRCh38 reference genome was built for each allele (Figure 2B). This adjusted reference contained the large structural variants of the DNA under study, based on the defined breakpoints. Then, the reads from both alleles were mapped once again, this time against the corresponding self-constructed and more representative reference sequence for each allele. After that, a first consensus sequence for each allele was deduced using the Medaka Consensus pipeline (v1.4.3) (Figure 2C). Subsequently, the consensus sequences for the two alleles were further optimized by mapping all the initially mapped reads to the GRCh38 target region. Reads that did not map unambiguously on one of the alleles were removed from the mapping data. Based on the newly mapped reads, the consensus sequences were finalized, and an accompanying probability file was generated using the Medaka Consensus pipeline (v1.4.3) (Figure 2D).

**Figure 2.**
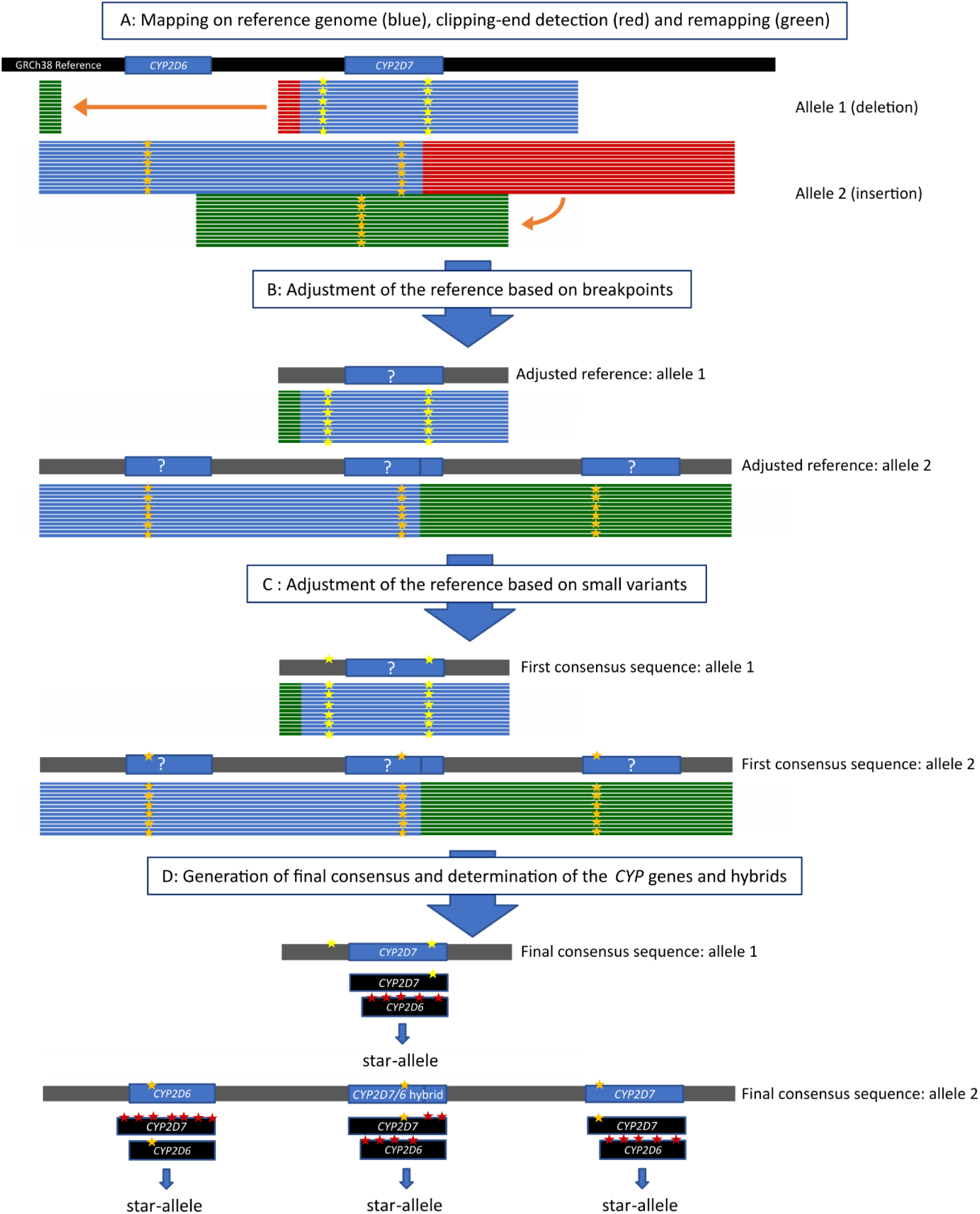
Workflow of the in-house developed CoLoRGen pipeline, which combines large structural and small variant calling. A: The basecalled reads are mapped against the human reference genome GRCh38 (black). Reads are split into the two alleles based on the small variants (yellow and orange stars). Clipping ends of the reads (red) are cut in-silico and mapped again to the reference genome (green). B: The reference is adapted based on the breakpoints of the clipping ends in the DNA under study (grey). Reads of alleles 1 and 2 are mapped against their respective adjusted reference sequence to create a first consensus sequence. C: The reference sequences are further adjusted by mapping all the previously mapped reads to end up with a final consensus sequence. D: The GRCh38 sequences of the *CYP2D6* and *CYP2D7* genes are mapped against the final consensus sequences. The GRCh38 gene or fragment containing the least mismatches (red stars) is assigned to the corresponding gene or fragment of the consensus sequence, resulting in the determination of the corresponding genes and hybrids. Finally, star-alleles can be assigned based on the determined variants.

Finally, the genes or hybrids in the consensus sequence were exactly identified based on their small variants (Figure 2D). For this purpose, the GRCh38 references of the *CYP2D6* and *CYP2D7* genes were mapped to the final consensus sequence of each allele, and mismatches between the consensus and the GRCh38 references were called using the Medaka Variant software (v1.4.3). The GRCh38 gene or fragment containing the least mismatches was assigned to the corresponding gene or fragment in the consensus sequence. Hybrids of *CYP2D6* and *CYP2D7* were reconstructed by concatenating these generated fragments, and a quality score was assigned to each small variant by considering the probability distribution on that exact position. By completing these steps, the number of copies of each gene and the exact composition of the hybrids were determined for each allele. After that, the star-alleles defined in PharmVar were assigned to the consensus alleles using a look-up algorithm based on the variants present in each gene (33). The star-allele or sub-allele most similar in terms of variants was assigned to the alleles of each sample.

The newly developed CoLoRGen pipeline was benchmarked using the NA12878 hybrid Genome in a Bottle Consortium (GIAB)-Platinum Genomes benchmark dataset described by Krushe *et al*. (34). VCF-files for the *CYP2D6* and *CYP2D7* genes of our data were separately compared with the benchmark dataset using the hap.py software (35). Visualizing the variants and verifying if they were correctly called and phased was done with in-house developed python scripts (36).

The sequencing data from the MinION run with NA12878 DNA was subsampled to determine the 16X minimum depth needed for reliable detection of small variants. Subsampling of the raw data was carried out using Seqtk (37). The CoLoRGen pipeline was run on each subsample. For each subsample, the depth of both genes was calculated, and the number of false- and true-positives was determined using in-house developed python scripts. In the subsampled datasets with depths below 16X on a gene, more than one false-positive variant popped up compared to the complete dataset. Therefore, a minimum depth of 16X on each allele of each gene was set as the lower limit for reliable small variant detection.

The CoLoRGen pipeline and the additional scripts are available via GitHub and can also be used for other genes when adapting the target gene regions and adding correct references for the star-alleles (36,38).

## Results and discussion

### Optimization of the nCATS experimental set-up

The *CYP2D6*-*CYP2D7* locus from the CEPH/UTAH pedigree 1463 sample NA12878 was first sequenced on a MinION flow cell to evaluate the cleavage and enrichment efficiency of the designed gRNAs, and to assess their off-target binding potential. Visualizing the mapped reads showed an additional cleavage place to the ones that were expected for the designed gRNAs. This additional cleavage place was due to off-target binding and cleavage of the RNP with gRNA9 (Figure S1). Therefore, gRNA9 was omitted in the subsequent sequencing runs. The eight remaining gRNAs were used to prepare a NA12878 Flongle library (NA_F_) to confirm the previous results. However, the selection of gRNAs still proved to be suboptimal, as the reads revealed the generation of smaller fragments. This was due to the high cleavage efficiency of the RNP with gRNA3, which as a result, created smaller fragments instead of increasing the depth on-target (Figure S2). Hence, gRNA3 was omitted in the subsequent sequencing runs as well. Furthermore, as almost no reads covering the complete *CYP2D6-CYP2D7* locus were present in the data from these preliminary sequencing runs, two pools with gRNAs were created for the subsequent runs. One pool did not contain the gRNA that cleaves between *CYP2D6* and *CYP2D7* to increase the number of reads covering the complete locus in the subsequent datasets.

### Enrichment of the *CYP2D6*-*CYP2D7* locus using nCATS or nCATS-AS

The enrichment efficiencies of both the nCATS-AS and the nCATS-only enrichment strategies were assessed during this study. For this purpose, the abovementioned nCATS enriched NA12878 library was sequenced on a MinION flowcell of which half of the pores were controlled by the AS software (NA_nCATS-AS_), and the other half of the pores were sequenced conventionally (NA_nCATS_). The NA_nCATS-AS_ data obtained an on-target depth of 128X, which was a 1.16 times increase compared to the NA_nCATS_ data (Table 1). After the preliminary sequencing runs with NA12878 libraries, two additional MinION runs were performed on libraries from extracted HG01990 (HG_nCATS-AS_ and HG_nCATS_) and GM19875 (GM_nCATS-AS_ and GM_nCATS_) DNA. The purpose of these runs was to evaluate if the enrichment strategies can generate correct *CYP2D6* and *CYP2D7* alleles for cell lines containing large structural variants. For these libraries, the two separate pools with the final selection of gRNAs were used. Furthermore, the same AS conditions as for the first MinION run were applied to additionally determine if AS exhibits added value for the enrichment of the *CYP2D6-CYP2D7* locus in these cell lines. The HG_nCATS-AS_ and HG_nCATS_ libraries reached an on-target depth of 25X and 30 X, respectively. Lower depths of 7X and 12X were obtained for the GM_nCATS-AS_ and GM_nCATS_, respectively (Table 1).

**Table 1.** General sequencing results of the nCATS-enriched NA12878, HG01990, and GM19785 libraries.

|  | NA12878 |  |  | HG01990 |  |  | GM19785 |  |  |
| --- | --- | --- | --- | --- | --- | --- | --- | --- | --- |
|  | nCATS-AS | nCATS | Combined | nCATS-AS | nCATS | Combined | nCATS-AS | nCATS | Combined |
| Throughput (MB) | 500 | 5 000 | **5,500** | 7 | 92 | **99** | 0.7 | 138 | **139** |
| Total reads | 588,959 | 2,213,701 | **2,802,660** | 1,470 | 11,066 | **12,536** | 771 | 18,778 | **19,549** |
| Reads on-target | 935 | 806 | **1,741** | 131 | 146 | **277** | 43 | 69 | **112** |
| Average target depth | 128X | 110X | **238X** | 25X | 30X | **55X** | 7X | 12X | **19X** |
| Percentage on-target (%) | 0.16* | 0.04* | **0.06** | 8.91 | 1.32 | **2.21** | 5.58 | 0.37 | **0.57** |

Each library was sequenced on one flow cell with half of the pores in AS mode, and half of the pores in uncontrolled mode. 'nCATS-AS' refers to the data of the pores in AS mode; 'nCATS' refers to the data generated by the uncontrolled, conventionally sequencing pores; 'combined' (values in bold) refers to the combined dataset containing both the positively selected reads from the AS pores and all the reads from the conventionally sequencing pores. \*: In this run, multiple *CYP*-genes were enriched with separate gRNA pools. Therefore, these on-target percentages should not be compared with the on-target percentages of the other runs.

The use of the AS software in addition to the nCATS enrichment did not consistently result in a higher on-target depth, but it did result in a considerably higher on-target percentage for all three cell lines (Table 1). However, as the vast majority of the strands were rejected by the software, the throughput generated by the AS controlled pores was also proportionally lower. Moreover, there were no more target strands encountered in the adaptive sequencing pores, as the rejected DNA strands were not removed from the flow cell, thus still hindering the accessibility of the pores. Overall, this resulted in approximately the same absolute number of on-target reads compared to the other pores, for which only nCATS-enrichment was used. Therefore, it can be concluded that the AS software does not conclusively offer sufficient additional benefit in this context. However, the advantages of adaptively sequencing certain specific strands have already been demonstrated in other contexts (27,39).

The enrichment efficiency of the nCATS strategy on itself was assessed as well. In their Cas9 targeted sequencing protocol, ONT mentions that a minimum target depth of 100X should be achievable (40). This depth was only obtained for the first MinION run in this study. All other runs reached a combined target depth of the AS-controlled and conventionally sequencing pores below 60X (Table 1). This value is expected to be influenced by two important factors that should be considered when determining the nCATS experimental set-up. The first factor is the number of gRNAs used for each target. ONT recommends using four gRNAs for regions smaller than 20 kb, two upstream of the target region and two downstream. Adding additional gRNAs at one side of the target region increases redundancy, so there is always at least one properly functioning gRNA in case of mutations in the recognition site of one of the other gRNAs at that position (26). As four gRNAs were designed upstream of *CYP2D6* and two downstream of *CYP2D7* in this study, this factor can be eliminated as a possible issue. The second factor to consider is the length of the input DNA. When the target region is longer than the average length of the input DNA, the depth drops towards the center part of the targeted region. Moreover, the target length increases when gene insertions or duplications are present, thereby complicating the achievement of sufficient depth even more. To increase the depth in the center of the targeted region, ONT advice is to follow the tiling approach, as described in their protocol (40). In the tiling approach, two pools of gRNAs are used. Each pool generates fragments that overlap with the fragments of the other pool. However, the downside of using the tiling approach is that fewer or no full-length reads of the gene construct are generated. To overcome this drawback, two different gRNA pools were composed in this study, one containing gRNAs that cut upstream and downstream the *CYP2D6-CYP2D7* locus, and another one also containing a gRNA cutting the DNA between the two genes. The input DNA was divided into two tubes, and each tube was incubated with a different gRNA pool to obtain reads covering the full *CYP2D6-CYP2D7* locus but also enrich the depth in the middle of the locus. Moreover, using a gRNA that cuts in the middle of the locus also aids in obtaining sufficient depth on *CYP2D6* for reliable variant calling. However, although these two factors were considered for our experimental set-up, the predetermined target depth was not obtained in this study.

Another factor influencing the obtained target depth is the percentage of on-target reads. PCR-free enrichment using nCATS generally resulted in a low percentage of on-target reads. Even after optimizing our customized pools of gRNAs for the *CYP2D6*-*CYP2D7* locus, a maximum on-target percentage of only 1.32% could be reached when this enrichment method was used without AS (Table 1). ONT reference samples comparable in length achieve an on-target percentage of 0.4% (26). Although our results are better, the obtained enrichment remains limited. Background DNA is assumed to be the main cause for this limited enrichment, as the number of off-target reads was only about 1%. The large amount of sequencable background DNA is probably due to the inefficiency of certain protocol steps or breakage of DNA strands when handling the DNA, making phosphorylated ends to which an adaptor can bind. Besides carefully executing the steps of the protocol, no other measurements could have been implemented to increase this percentage. Logically, this low obtained percentage of on-target reads on its turn resulted in a low depth on target. However, this is not the only factor inherent to the nCATS protocol that influences the maximum obtainable target depth.

The overall throughput of the sequencing run also plays an important role in obtaining sufficient target depth. The nCATS protocol generated low throughputs for all three DNA samples (Table 1). This is caused by the presence of non-adaptor-ligated DNA strands in the flow cell, as these are not removed during the library preparation. These DNA strands are assumed to spatially block the pores, thereby hindering the sequencing of the adaptor-ligated DNA strands and causing a very low pore occupancy. The low target depth ensuing from the background and non-adaptor-ligated DNA strands comprises one of the main disadvantages of the nCATS enrichment method in the pharmacogenetics context. It implies that one flow cell per patient is needed to get enough depth on the targeted region(s), resulting in a high sequencing cost that hinders the implementation of the proposed assay in practice. Optimizing the nCATS protocol by incorporating an additional purification step for the adaptor-ligated strands might solve this issue and increase the on-target depth, allowing multiple samples to be sequenced on one flow cell. The establishment of a purification step compatible with the nCATS-protocol constitutes the follow-up research to this paper.

### SNV and INDEL calling performance on reference NA12878 DNA

The small variant calling performance of the nCATS enrichment strategy combined with the CoLoRGen analysis pipeline was assessed using the NA12878 library, as only for this DNA a truth set containing all small variants is available in the literature (34). For this purpose, the NA_combined_ dataset was used, combining the nCATS-AS and the nCATS reads, as the only difference between these reads is the specific pore on the same flow cell it was sequenced on. The truth set composed by Krusche *et al*. (34) contains 11 SNVs and 1 INDEL in the *CYP2D6* gene, and 26 SNVs and 1 INDEL in the *CYP2D7* gene (Figure 3). All 11 and 26 SNVs in *CYP2D6* and *CYP2D7*, respectively, were also called and phased in the NA_combined_ dataset (Figure 3). However, two additional, supposedly false-positive SNVs were called in *CYP2D6*, and five in *CYP2D7*. As for the INDELs, only the deletion in *CYP2D6* was called and phased correctly. The insertion in *CYP2D7* remained undetected, but four additional deletions were detected in the NA_combined_ consensus of *CYP2D7* instead. Remarkably, all supposedly false-positive SNVs and INDELs in both genes were assigned to the same allele after phasing. This raises the question as to whether the NA12878 reference by Krusche *et al*. is incorrect, and consequently the false-positive variants are actually present in the NA12878 DNA. Additional results and discussions on this can be found in the sections below.

**Figure 3.**
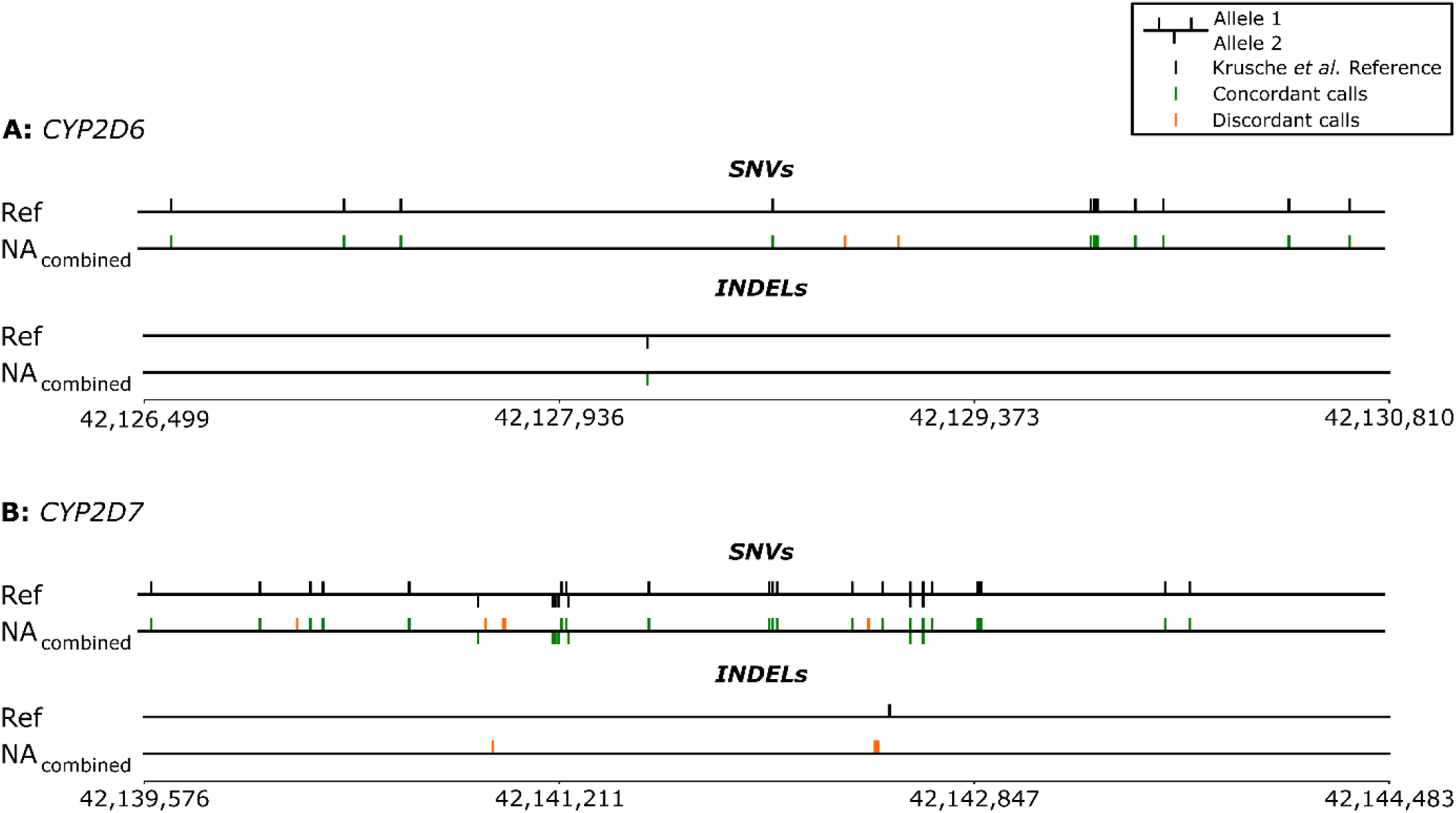
Representation of the called and phased small variants (SNVs and INDELs) in the *CYP2D6* and *CYP2D7* genes of the NA_combined_ library. The truth set composed by Krusche *et al*. (34) was used as reference (Ref). Green lines represent concordant calls (true-positives compared to the truth set), which are correctly called and phased variants compared to the reference; orange lines represent discordant calls (false-positives compared to the truth set). Note: multiple variants next to each other are visually represented by thicker lines.

### Comprehensive genotyping of the NA12878 *CYP2D6-CYP2D7* locus by the CoLoRGen pipeline

The CoLorGen pipeline detected a structural variant in addition to the small variants in the NA12878 DNA. Based on all the detected variants, CoLoRGen assigned the *CYP2D6* *3/*4+*68 star-alleles to the NA_combined_ dataset, of which the *68 allele represents a *CYP2D6-CYP2D7* hybrid insertion (Figure 4). The high obtained on-target depth of 238X implicates that the detection of this hybrid cannot be attributed to nanopore sequencing errors or an artifact of the analysis pipeline. However, no large structural variants have been identified for the *CYP2D6-CYP2D7* locus in the NA12878 hybrid benchmark of Krusche *et al*. (34). Accordingly, the Get-RM studies did not unambiguously assign a structural variant to the NA12878 DNA (15,16). In these Get-RM studies, several testing laboratories conducted different assays, but only when TaqMan-based genotyping was combined with CNV and structural variant detection using quantitative multiplex PCR and LR-PCR validation, the presence of the *68 hybrid could be detected (15). Therefore, the *68 allele was not included with 100% certainty in the reported consensus star-allele classification (15). In accordance with our results, a more recently published article also reported the statistical inference of the *68 allele in NA12878 whole-genome sequencing (WGS) data when using the Cyrius analysis tool (41). As the *68 hybrid has been inferred in the NA12878 DNA multiple times in literature, it can be concluded that this structural variant is effectively present and was thus correctly identified by the CoLoRGen pipeline.

Furthermore, it was noted that the hybrid was phased to the same allele as all the supposedly false-positive SNVs and INDELs. As the hybrid was not included in the NA12878 reference provided by Krusche *et al*. (34), other variants may also be incorrectly identified in that reference due to the incorrect mapping of the reads originating from the *CYP2D6-CYP2D7* hybrid on the *CYP2D6* or *CYP2D7* gene. This can be substantiated with the fact that the reference data set for the NA12878 DNA is mainly constructed based on Illumina short-read sequencing data and older versions of the long-read sequencing technologies, which are more prone to generating inaccurate sequences for complex loci as *CYP2D6-CYP2D7* (42,43). These results indicate that the NA12878 references might be outdated and not entirely accurate, and highlight the advantage of the nCATS enrichment strategy combined with the CoLoRGen pipeline, which can simultaneously detect large structural and small variants.

Some other published assays also correctly determine the presence of the *CYP2D6-CYP2D7* *68 allele. However, our nCATS-CoLoRGen assay has added value by providing the complete allele sequences spanning the entire *CYP2D6-CYP2D7* locus, including the exact structural variant sequence. None of the reported assays provide this comprehensive information to the best of our knowledge. LR-PCR could be used as an alternative enrichment strategy, but is mostly only able to target *CYP2D6* (20). Larger regions, including *CYP2D6, CYP2D7*, and possible deletions, duplications, and hybrids, are difficult to cover with LR-PCR since the probability of getting chimeric molecules increases with the length of a PCR amplicon (23). TaqMan genotyping combined with quantitative multiplex PCR and LR-PCR validation, or short-read sequencing combined with the statistical modeling and counting Cyrius tool are genotyping approaches that could detect the presence of the *68 hybrid (15,41). Nevertheless, these assays also do not directly provide the allele-specific sequence of the locus, but are instead used to classify the *CYP2D6* locus into a predefined set of star-alleles. However, the current classification of CYP2D6 enzyme activities based on the star-allele gene definitions has proven to be a suboptimal predictor for enzyme activity (44). More recent research by Van der Lee *et al*. (45) supported this by confirming that building a predictive model based on the complete *CYP2D6* gene sequence gives better predictive values for the gene function than a model built solely based on the star-alleles. By generating complete consensus sequence, CoLoRGen can phase additional mutations, thereby allowing a more accurate gene function predictions.

### Validation of genotyping performance using two additional cell lines

The DNA of two additional cell lines, HG01190 and GM19785, was used to verify the structural variant detection performance of the nCATS-CoLoRGen pipeline. The HG01190 cell line contains two major structural variants (15). One allele has a complete deletion of the *CYP2D6* gene, referred to as the *5 allele. The other, *4+*68 allele, contains a duplication, defined as a hybrid between *CYP2D7* and *CYP2D6* (Figure 4). The HG_combined_ dataset contained 37 reads that covered the breakpoints of the 12,152 basepair-long deletion between positions 42,123,191 and 42,135,343 (Figure S3). Additionally, a 13,680 basepair-long duplication of the region between positions 42,145,873 and 42,132,193 was discovered in six reads. As more than three reads were covering the breakpoints of the large structural variants, the deletion and insertion were considered to be detected correctly. Subsequently, detection of the small variants was used to exactly identify *CYP2D6, CYP2D7*, or possible hybrids. The minimum 16X depth for reliable small variant calling was obtained on all detected gene copies except on the insertion of allele 2. Nevertheless, the cell line was correctly identified as the *5/*4+*68 genotype by our CoLoRGen pipeline (Figure 4).

**Figure 4.**
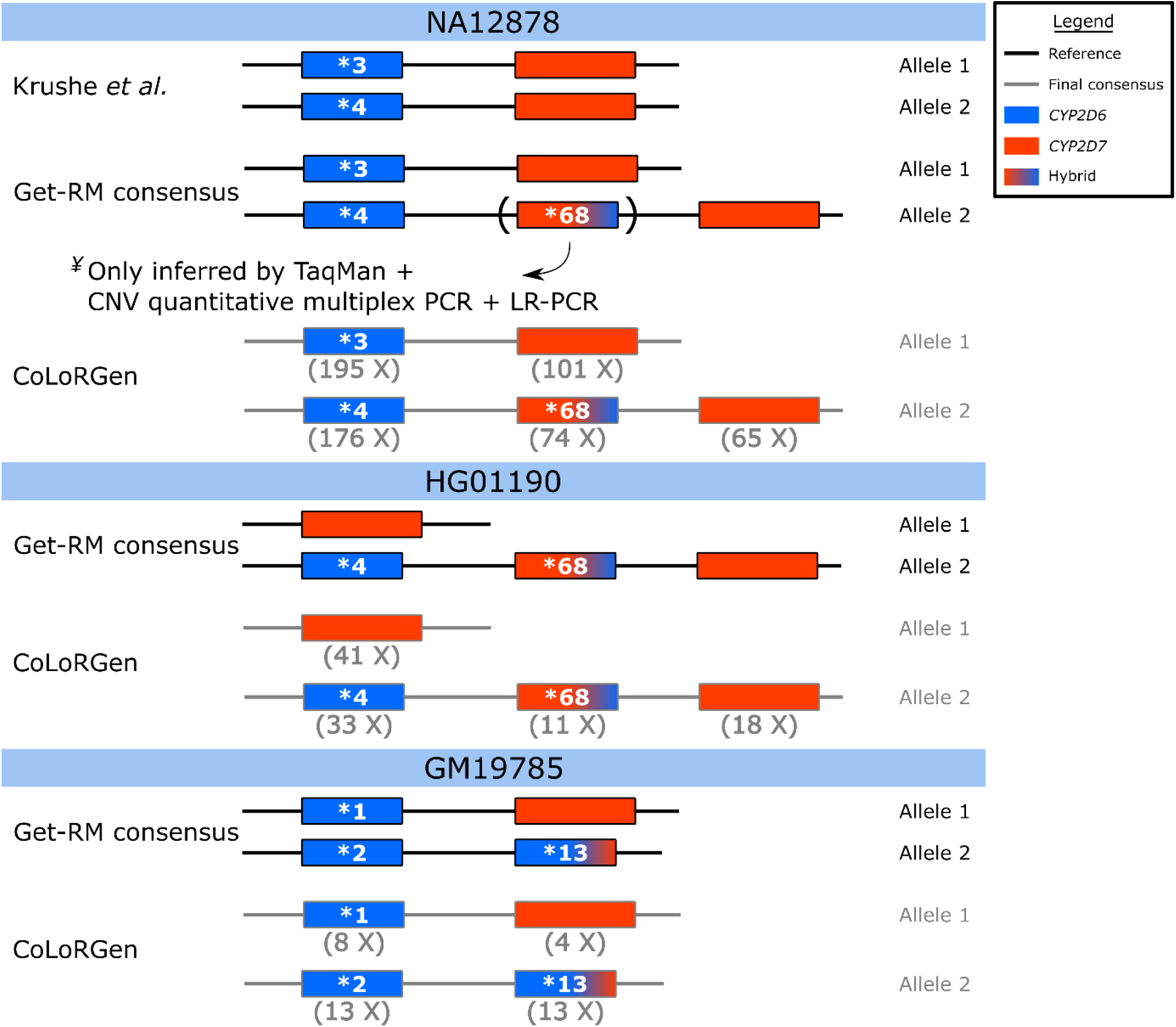
Star-alleles in literature references and star-alleles assigned by the CoLoRGen pipeline. Reference star-alleles were obtained from Krushe *et al*. (34) and the Get-RM studies (15,16). The depths mentioned below the genes are the generated average depths on that position of the locus. ^¥^The *68 allele was only detected when TaqMan-based genotyping was combined with CNV and structural variant detection using quantitative multiplex PCR and LR-PCR validation. Therefore, the Get-RM consensus star-allele only mentions the *68 allele in brackets. Note: even when depths below the minimal 16X depth for reliable small variant calling were obtained, correct star-alleles could be assigned.

The GM19785 cell line consists of a *1 allele, without any structural variants, and a *2+*13 allele, containing one *CYP2D6* copy and a *CYP2D6-CYP2D7* hybrid (Figure 4) (15). The hybrid replaces the *CYP2D7* gene in this allele, which implies that there is no difference in the number of gene copies, but only a difference in the DNA sequence on the exact position where *CYP2D7* is normally located. However, the *CYP2D6-CYP2D7* hybrid can map on *CYP2D7* due to their highly similar sequences. Therefore, the CoLoRGen pipeline can only detect this structural variant based on the small variants in the gene sequence, and not based on mapped reads with clipping ends. Although insufficient target depths below 16X were reached on both alleles of the GM_combined_ dataset, our CoLoRGen pipeline could assign the correct *1/*2+*13 genotype to the GM19785 DNA (Figure 4).

The exact sequence between the *CYP2D6* gene and the *CYP2D6-CYP2D7* hybrid could not be determined for the GM19875 cell line, as no reads covering the whole target region were generated. This is due to the presence of a part of the *CYP2D6* sequence at the start of the *CYP2D6-CYP2D7* hybrid, which introduced an additional recognition site for gRNA2 that is normally only present upstream of the *CYP2D6* gene locus. The additional recognition site was visible in the mapped reads, as all the reads were cut in the middle at the same cleavage site (Figure S4). This problem might arise when hybrids are present in the target sequence, but can be circumvented by designing gRNAs located further away from the target gene. However, the further a gRNA is located from the target, the lower the obtained on-target depth will be. This is a trade-off that should be taken into account when designing optimal gRNAs.

### In-depth discussion of the generated consensus sequences

Although the CoLoRGen pipeline could assign the correct star-alleles to the studied samples, a further in-depth analysis revealed the presence of additional small variants in the final consensus sequences, besides the variants that were assigned to a specific star-allele. Most of these additional variants are present in several sub-allele definitions, thereby confirming the correct assignment of the star-allele. Nevertheless, some additional or lacking variants were often observed in our data compared to the exact sub-allele definition. In the *4 allele of the NA_combined_ and HG_combined_ libraries, 12 additional variants were detected, which were exactly the same for both samples. These variants are all included in several defined sub-alleles, but these sub-alleles contain other variants in addition. In the *1 allele of the GM_combined_ data, two additional deletions were called. One of them was situated in an intron, and the other in an exon region. Both additional deletions were located in homopolymeric regions. The *2 allele of the GM_combined_ data contained 13 additional variants denoted in several *2 sub-allele definitions. Two other additional variants in our data are not defined in the star- or sub-allele database (5) and were both located in exon regions. One of these variants was located in a homopolymeric region. The other variant was not located in a homopolymeric region but represents a synonymous mutation. Therefore, it does not impact the resulting amino acid sequence (Figure S5).

The four additionally detected variants that were not present in the star- or sub-allele definitions were all from the GM_combined_ dataset, which had insufficient depths for reliable small variant calling (Figure 4). Moreover, three out of these four variants were INDELs located in homopolymeric regions, which are notoriously error-prone regions in ONT sequencing. Therefore, these additionally called variants are probably due to nanopore sequencing errors. The R10.3 flow cell, which has a better performance in homopolymeric regions, was available at the time of writing and is supposed to overcome this problem. However, we decided not to sequence this library on an R10.3 flow cell, as more random errors seem to occur when using this type of flow cell, and R9.4 flow cells still prove to provide better genotyping results (46,47). Nevertheless, efforts are still made by ONT to improve the consensus accuracy of homopolymer regions, which holds promising perspectives for obtaining better results in the future. Another possible explanation for the additional detected variants can be found in the star-allele nomenclature itself. These definitions are intrinsically not comprehensive, as only variants based on microarrays and known effects on the enzyme level are considered in their definitions. Non-coding variants were only considered for recently added star alleles (6). Even though this nomenclature is not optimal in our context of defining complete alleles, the star-allele definitions were used to benchmark our results as no other definitions were yet available at the time of writing. However, a new and more comprehensive system to document gene sequences in the pharmacogenetic field should be a general objective for the future, as the current nomenclature is somewhat outdated.

### Variant calling performance of CoLoRGen pipeline *versus* state-of-the-art variant callers

To determine the added value of the newly developed CoLoRGen pipeline, a comparison was made with state-of-the-art variant callers. However, existing small variant detection tools cannot detect large structural variants, and, accordingly, large structural variant detection tools cannot detect small variants. Therefore, separate comparisons were made for the detection of small SNVs and INDELs on the one hand, and large structural variants on the other hand.

First, the NA_combined_ dataset was analyzed with the Medaka Variant pipeline to compare the SNV and INDEL calling performance of the CoLoRGen pipeline with the state-of-the-art small variant caller for nanopore sequencing data (31). Although CoLoRGen did not call all SNVs and INDELs correctly, the results were comparable with the results generated by the Medaka Variant pipeline (Table S2). The called SNVs and INDELs that differed between both variant callers were either located in a homopolymeric region or in a region where CoLoRGen detected a hybrid insertion. Homopolymeric regions are a known cause for nanopore sequencing errors and are therefore likely to be responsible for the generation of false-positive small variants (48). Furthermore, regions containing large structural variants, such as hybrid insertions, cannot be detected by the Medaka Variant pipeline. Consequently, reads originating from the hybrid are incorrectly mapped on *CYP2D6* or *CYP2D7* when using the Medaka Variant pipeline, giving rise to more called SNVs and INDELs. However, as the small differences in results between both pipelines can be explained by these two causes, our CoLoRGen pipeline proved to perform adequately for calling SNVs and INDELs in complex genes such as *CYP2D6*. Moreover, as the CoLoRGen pipeline combines both large structural and small variant calling, it can generate a more comprehensive genotype in comparison with the Medaka Variant pipeline.

Second, the NA_combined_, HG_combined_, and GM_combined_ datasets were also analyzed with the existing large structural variant detection tools NanoVar (30), Sniffles (29), and SVIM (28) to compare the large structural variant calling performance. None of these tools was able to reliably elucidate all the large structural variants in the complex *CYP2D6-CYP2D7* locus of the cell lines used in this study (Table S3). Additionally, the output of these tools is not easily interpreted. Therefore, the CoLoRGen tool outperformed these tools as well in terms of generating a correct and comprehensive genotype of the complex *CYP2D6-CYP2D7* locus. When aiming for a suitable pharmacogenetic assay to use in clinical practice in the future, a comprehensive and straightforward data analysis tool is of major importance, hence the usefulness of this developed comprehensive CoLoRGen pipeline.

## Conclusion

In this study, the enrichment efficiencies of the nCATS and the nCATS-AS strategies were assessed on the *CYP2D6-CYP2D7* locus in aiming to develop an assay that can accurately genotype complex pharmacogenes. In addition, we developed and evaluated CoLoRGen, a new and more comprehensive analysis pipeline to simultaneously detect both large structural and small variants. The nCATS-CoLoRGen assay resulted in the assignment of correct star-alleles to the *CYP2D6* gene and *CYP2D6-CYP2D7* hybrid in 3 cell lines containing complex gene structures. Moreover, the CoLoRGen pipeline also generated a complete consensus sequence of the genes, thereby demonstrating the presence of *CYP2D6-CYP2D7* large structural variants and smaller SNVs and INDELs that go undetected by other current methods. Our results provide direct evidence that the *CYP2D6* genotype of the NA12878 DNA should include the *CYP2D6-CYP2D7* *68 hybrid and several additional SNVs compared to existing references (15,16,34). However, the implementation of this assay in practice is hampered by the fact that both the nCATS and nCATS-AS strategies led to a low percentage of on-target reads, resulting in low on-target sequencing depths. Further optimization of the nCATS enrichment strategy is thus worth considering for following research, as the usefulness of a long-read PCR-free enrichment strategy in combination with the CoLoRGen pipeline for accurate gene function predictions has been demonstrated in this study.

## Availability of data and materials

The datasets generated and analyzed during the current study are available as BioProject, PRJNA796180

The CoLoRGen pipeline and other used code are available at GitHub: https://github.com/laurentijntilleman/CoLoRGen

## Competing interests

The authors declare that they have no competing interests

## Funding

KD and this research are supported by the Special Research Fund (Bijzonder Onderzoeksfonds, BOF, University Ghent, BOF21/DOC/042)

## Authors’ contributions

KR: Conceptualization, Methodology, Investigation, Writing – Original Draft, Visualization; LT: Conceptualization, Methodology, Software, Formal Analysis, Investigation, Data Curation, Writing – Original Draft, Visualization; KD: Investigation, Writing – Review & Editing; OT: Methodology, Writing – Review & Editing; DD: Writing – Review & Editing, Funding Acquisition; FV: Conceptualization, Methodology, Writing – Review & Editing, Supervision, Funding Acquisition

## Supplemental material

**Figure S1.**
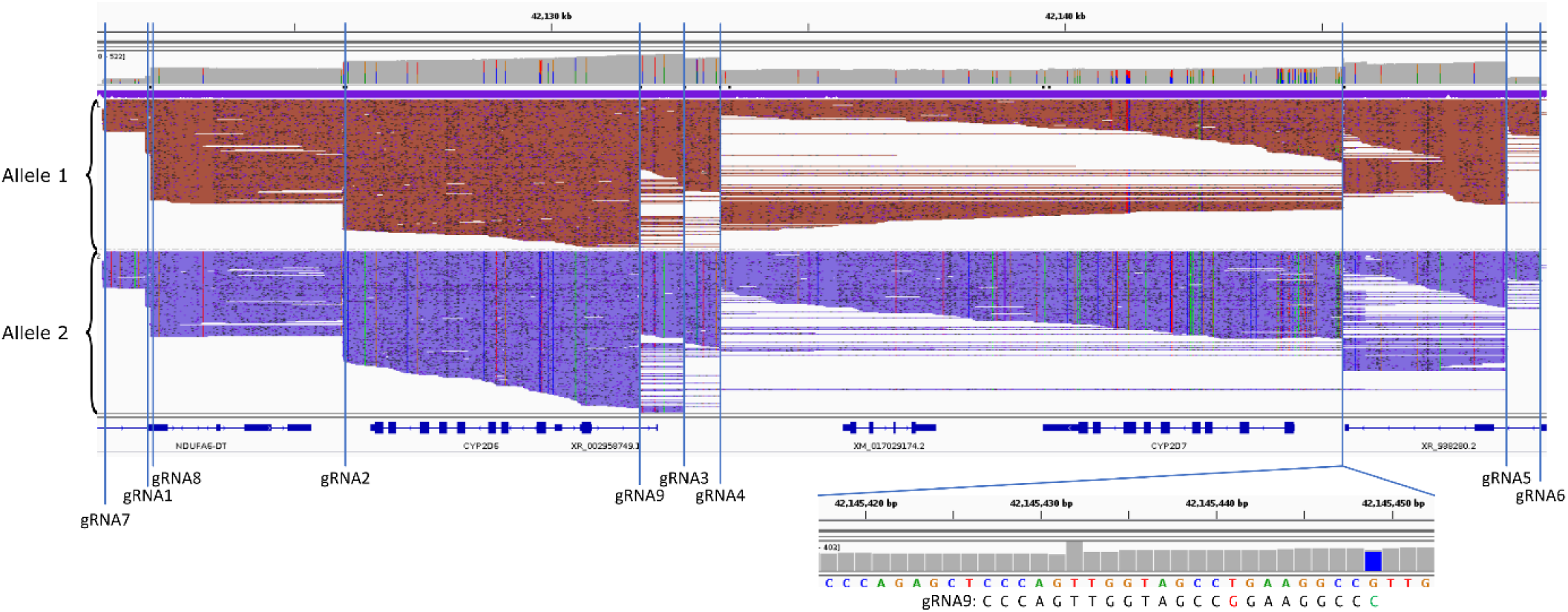
Mapped reads of the NA12878 DNA sequenced on a MinION flow cell. The positions of the gRNAs are indicated with vertical lines. Reads are split by allele. The position where gRNA9 binds off-target is zoomed in. This recognition site shows one mismatch (red) and one mutation (green).

**Figure S2.**
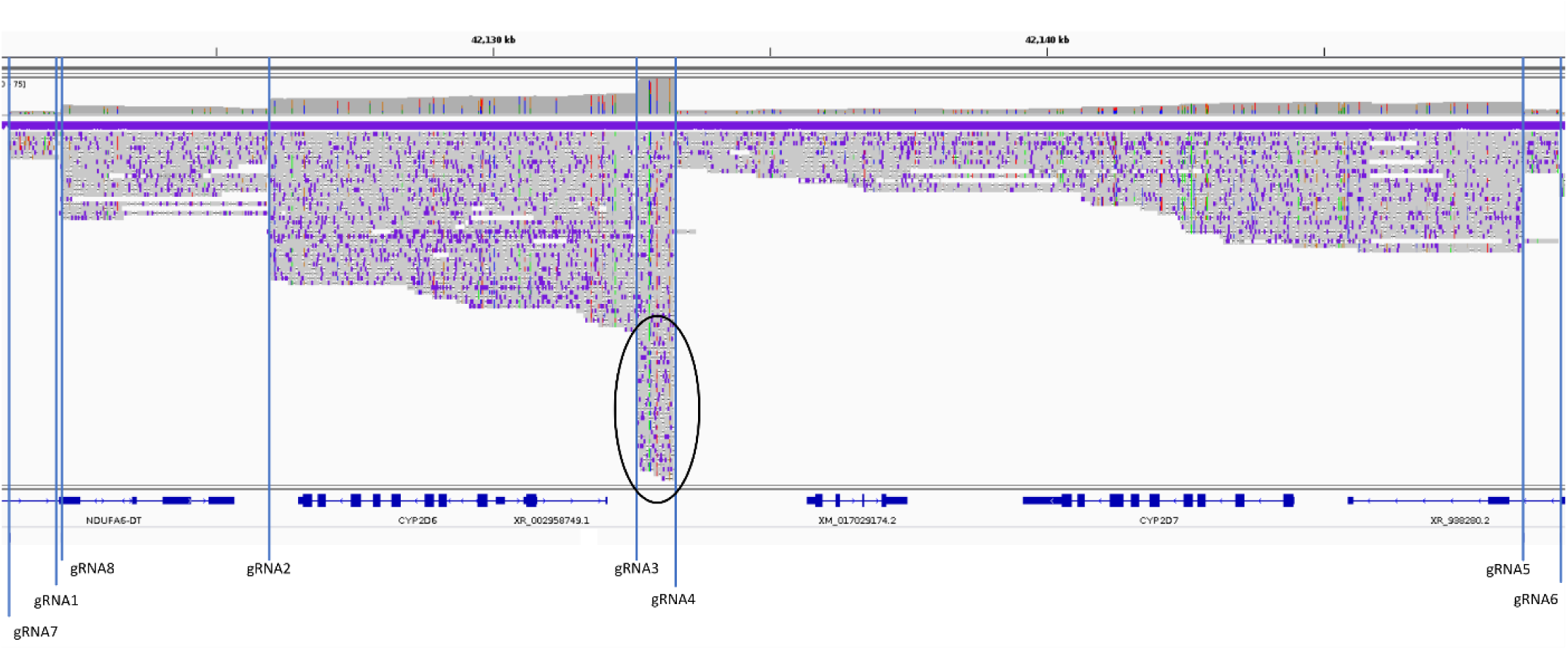
Mapped reads of the NA12878 DNA sequenced on a Flongle flow cell. The positions of the gRNAs are indicated with vertical lines. gRNA3 cut reads generated by gRNA4, causing a lower depth on *CYP2D6*.

**Figure S3.**
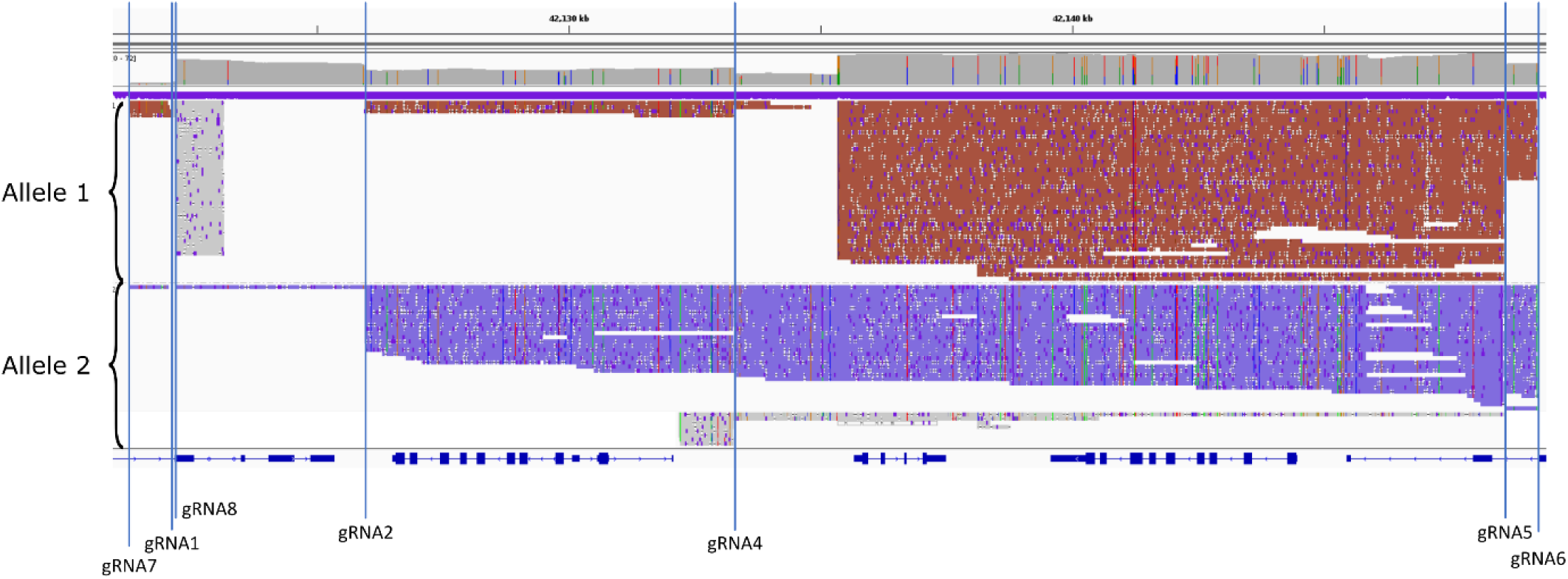
Mapped reads of the HG01190 DNA sequenced on a MinION flow cell. The HG_combined_ dataset was used to generate this figure, which is the dataset containing both the positively selected reads from the AS pores and all the reads from the conventionally sequencing pores. The positions of the gRNAs are indicated with vertical lines. Reads are split by allele, and gray reads are clipping ends that were cut in-silico and mapped separately.

**Figure S4.**
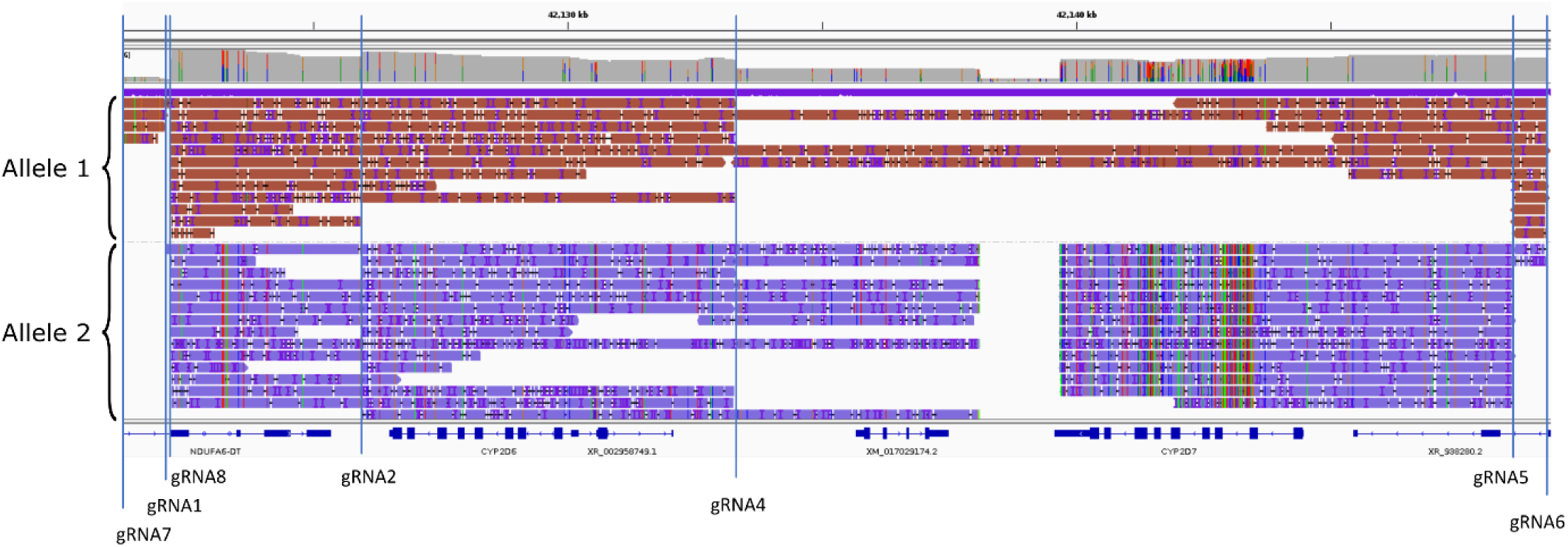
Mapped reads of the GM19785 DNA sequenced on a MinION flow cell. The GM_combined_ dataset was used to generate this figure, which is the dataset containing both the positively selected reads from the AS pores and all the reads from the conventionally sequencing pores. The positions of the gRNAs are indicated with vertical lines. Reads are split by allele.

**Figure S5.**
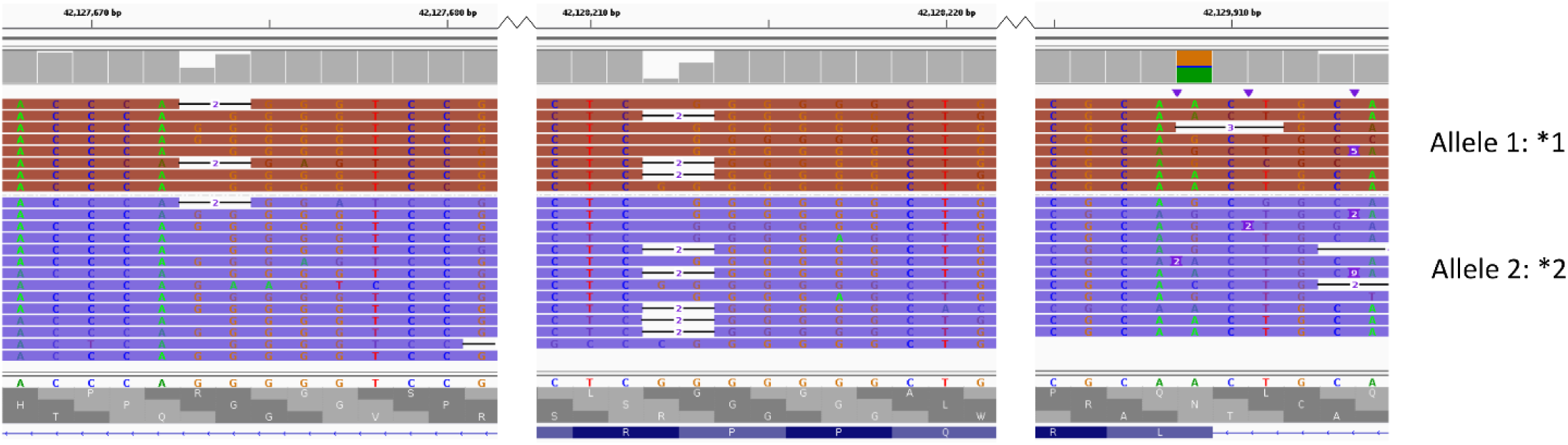
CoLoRGen detected four additional small variants in the GM19785 cell line that are not present in the sub-allele definitions. The three deletions were located in homopolymeric regions and the SNV is a silent mutation.

**Table S1.**
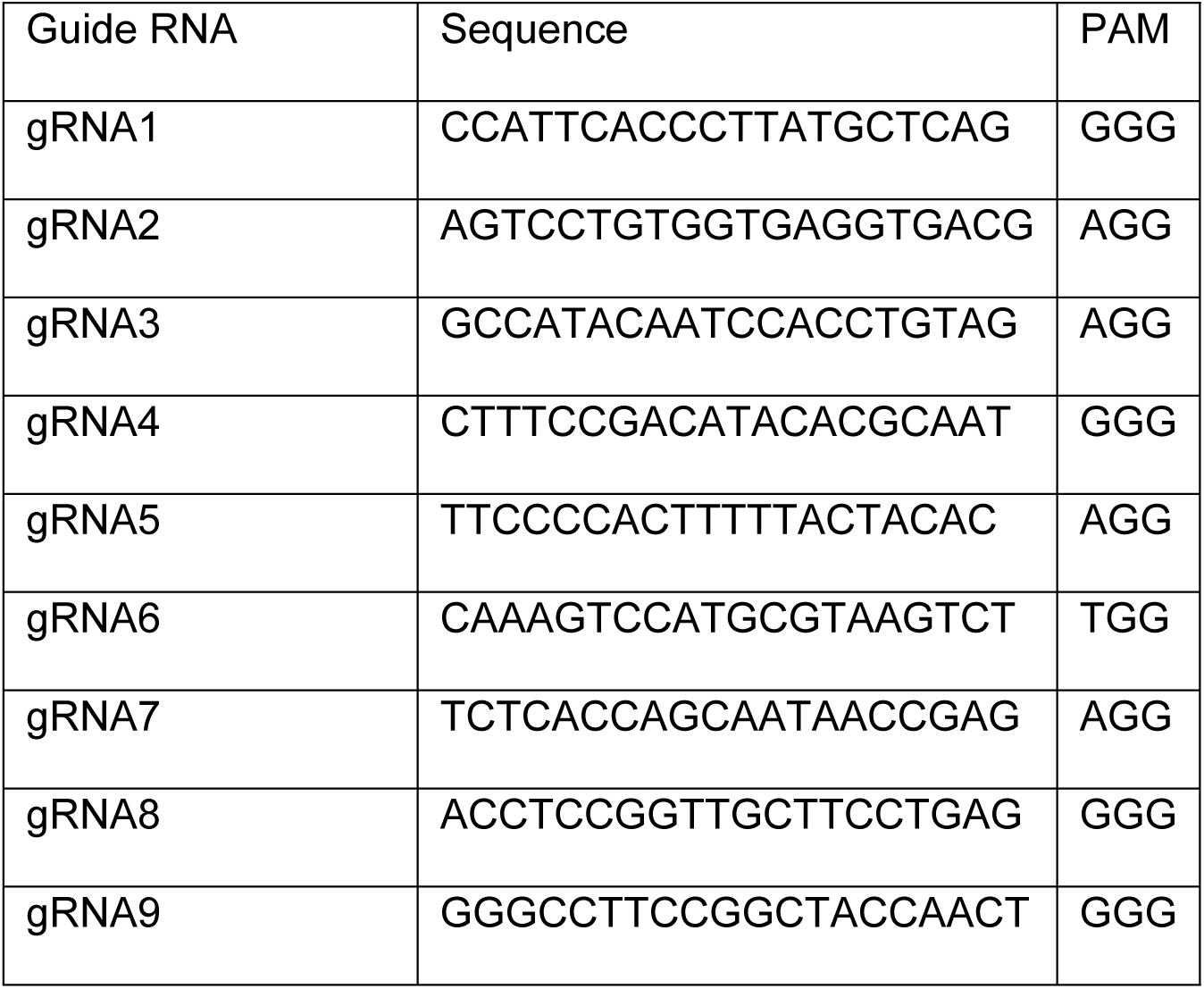
Overview of the used guide RNAs (gRNAs).

**Table S2.**
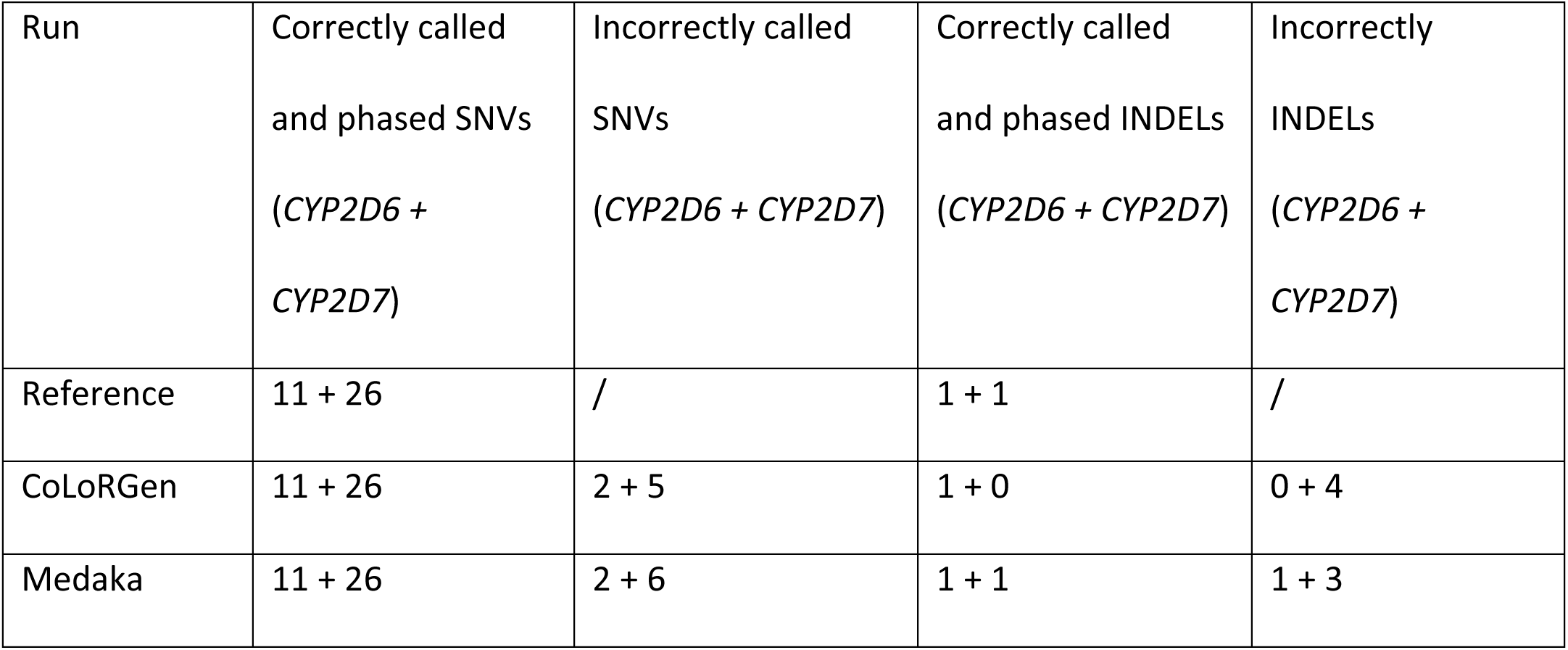
Comparison of small SNV and INDEL variant detection of the Medaka Variant pipeline and the new CoLoRGen tool in the NA12878 DNA sample. Reference: Krusche *et al*. (34).

| Run | Correctly called<br>and phased SNVs<br>(*CYP2D6* +<br>*CYP2D7*) | Incorrectly called<br>SNVs<br>(*CYP2D6* + *CYP2D7*) | Correctly called<br>and phased INDELs<br>(*CYP2D6* + *CYP2D7*) | Incorrectly<br>INDELs<br>(*CYP2D6* +<br>*CYP2D7*) |
| --- | --- | --- | --- | --- |
| Reference | 11 + 26 | / | 1 + 1 | / |
| CoLoRGen | 11 + 26 | 2 + 5 | 1 + 0 | 0 + 4 |
| Medaka | 11 + 26 | 2 + 6 | 1 + 1 | 1 + 3 |

**Table S3.** Comparison of *s*tructural variant detection of different state-of-the-art structural variant tools and the new CoLoRGen tool in the NA12878, HG01190 and GM19785 DNA samples. For each tool the number of deletions and insertions are given. Between parentheses the length of each variant is given. Green: correctly detected structural variant; red: incorrectly detected structural variant; orange: multiple overlapping structural variants are detected although only one variant is present in the reference. Reference: Get-RM studies (15,16). †: the found regions show overlap.

|  | NA12878 |  | HG01190 |  | GM19785 |  |
| --- | --- | --- | --- | --- | --- | --- |
|  | deletion | insertion | deletion | insertion | deletion | insertion |
| Reference | / | *68 | *5 | *68 | / | / |
| CoLoRGen | / | 1 (13,680 bp) | 1 (12,152 bp) | 1 (13,680 bp) | / | / |
| NanoVar<br>(PASS) | / | / | / | 1 (13,838 bp) | / | / |
| Sniffles<br>(PASS) | 2 (12,282 bp,<br>12,152 bp) † | 3 (12,154 bp,<br>13,708 bp,<br>13,659 bp) † | 2 (12,454 bp,<br>12,155 bp) † | 1 (1,006 bp) | 1 (13,656 bp) | / |
| SVIM (QUAL<br>≥3, PASS) | / | 2 (13,638 bp,<br>13,613 bp) † | / | 1 (13,424 bp) | 2 (13,696 bp,<br>13,663 bp) † | / |

## Notes

### Competing Interest Statement

The authors have declared no competing interest.

